# Road and landscape-context impacts on bird pollination in a Mediterranean-type shrubland of the southeastern Cape Floristic Region

**DOI:** 10.1101/815324

**Authors:** B. Adriaan Grobler, Eileen. E. Campbell

**Author notes:** Corresponding author; Botany Department, PO Box 77000, Nelson Mandela University, Port Elizabeth 6031, South Africa.

## Abstract

Road verges can provide important habitats for plants, especially in transformed landscapes. However, roads and their associated traffic have several adverse impacts on ecosystems that can disrupt vital ecological processes, including pollination. In transformed landscapes, road effects on pollination might be complemented by impacts of large-scale habitat modification. In these landscapes, road verge populations of plants that rely on pollinators for pollen transfer could thus be at risk of pollination failure. This study investigates the pollination of a reseeding, bird-pollinated shrub, *Erica glandulosa*, in road verges of a fragmented and transformed rural landscape in the southeastern Cape Floristic Region. We test for road impacts on pollination by comparing number of ruptured anther rings—a proxy for pollination—in fynbos vegetation fragments at different distances from the road (0–10, 20–30 and 40–50 m). We also test whether different land-cover types (intact fynbos, alien thickets and rangelands/pastures) next to road verges influence the number of ruptured anther rings. After controlling for robbing rate and plant density, fewer flowers were pollinated near the road than farther away, and fewer flowers were pollinated where road verges occurred next to alien thickets or pastures/rangelands compared to intact fynbos. However, bird pollination was not excluded in road verges: on average, ca. 20–30% of flowers were still visited by birds near the road. These findings potentially call into question the suitability of road verges as refugia for seed-dependent, bird-pollinated plant species in transformed landscapes.

## 1. Introduction

Road verges have been highlighted as important habitats for plants, especially in transformed landscapes where large expanses of natural vegetation have been converted to alternate forms of land use (Arenas et al., 2017; Cale and Hobbs, 1991; Deckers et al., 2004; Leigh and Briggs, 1992; Lewis, 1991; Nicolson, 2010; O’Farrell and Milton, 2006). Although roads and verges can facilitate the spread of alien invasive plants (Ernst, 1998; Gelbard and Belnap, 2003; Hayasaka et al., 2012; Kalwij et al., 2008; Pauchard and Alaback, 2004; Rahlao et al., 2010; Tyser and Worley, 1992), where they are adequately managed, road verges may provide refuge to native plants in landscapes that are otherwise transformed by alien plant invasions. Thus, road verge habitats have become increasingly recognised for their contributions to nature conservation and the maintenance of landscape connectivity (Byrne et al., 2007; Hogbin et al., 1998; Pauwels and Gulinck, 2000; Van Rossum et al., 2004).

Despite these benefits, roads and their associated traffic have several adverse impacts on ecosystems (Coffin, 2007; Forman and Alexander, 1998; Spellerberg, 1998; Trombulak and Frissell, 2000) that can undermine the conservation value of road verge habitats. Roads modify animal behaviour through home range shifts and by altering reproductive success, movement patterns, escape responses, and physiological states (Trombulak and Frissell, 2000). Through this modified behaviour, roads and traffic exert population-level effects by reducing the density or size of animal populations near roads (Fahrig and Rytwinski, 2009; Rytwinski and Fahrig, 2015). In turn, these effects can lead to fragmentation of animal populations (Fahrig et al., 1995; Huijser and Bergers, 2000; Keller and Largiader, 2003), but, more importantly in the context of this study, could also disrupt vital ecological processes facilitated by animals, such as pollination.

Most flowering plant species—an estimated 87.5% (Ollerton et al., 2011)—depend on animals, especially insects, birds, bats, and small mammals, for pollination (Kearns et al., 1998). Road verge populations of plants that rely on pollinators for pollen transfer could thus be at risk of pollination failure, subsequent reproductive failure, and ultimately local extinction (Bond, 1994; Pauw, 2007; Trombulak and Frissell, 2000; Wilcock and Neiland, 2002). Even though certain plants may be able to compensate for insufficient pollination in the short-term, for example through clonal propagation (Pauw and Bond, 2011; Pauw and Hawkins, 2011) or vegetative sprouting (Bond and Midgley, 2001), it is debatable whether such mechanisms can facilitate the long-term persistence of species (Bond, 1994), especially considering predicted climate change.

In the fynbos ecosystems of South Africa’s Cape Floristic Region (CFR), plant species most at risk of this road-related impact are those that rely on birds for pollination, and whose regeneration depend on seeds. Obligate reseeding plants constitute about half the Cape flora, can be dominant components in fynbos communities, and are well represented in the Ericaceae and Proteaceae (Le Maitre and Midgley, 1992). Ornithophilous members of the Ericaceae and Proteaceae also typically dominate late seral communities in fynbos vegetation (Van Wilgen, 1981). The CFR is comparatively rich in bird-pollinated plants, with roughly 75% of southern Africa’s ornithophilous plant species occurring in the Cape, where they constitute ca. 4% (around 320 species) of the regional flora (Rebelo, 1987). Pollination by sunbirds (Nectariniidae) in particular is more extensively developed in the CFR than elsewhere in Africa (Manning and Goldblatt, 2012), with more than 44 plant species being exclusively pollinated by a single bird species, the long-billed malachite sunbird (*Nectarinia famosa*) (Anderson et al., 2005; Geerts and Pauw, 2009). Shorter-billed sunbirds, like the orange-breasted sunbird (*Anthobaphes violacea*) and southern double-collared sunbird (*Cinnyris chalybea*), pollinate over 66 species of plants in the Cape (Geerts and Pauw, 2009; Rebelo et al., 1985, 1984), while the cape sugarbird (*Promerops cafer*, Promeropidae) pollinates at least 20 plant species, typically members of the Proteaceae (Johnson, 2010). This guild of nectarivorous birds thus plays a central role in the reproduction of a significant portion of the Cape flora.

Whereas the direct impacts of roads and traffic on avian populations have received much attention, especially habitat fragmentation and road kill, few studies have investigated the behavioural response of birds to roads (Kociolek et al., 2011; Rytwinski and Fahrig, 2015). Kociolek et al. (2011) found that birds avoid roads due to the physical barrier roads present to movement, and due to artificial light, edge effects, and traffic noise. However, response to roads may be guild- or species-specific (Kuitunen et al., 1998; Laurance et al., 2004; Reijnen et al., 1997, 1996), with certain birds, mostly raptors and scavengers, even being attracted to roads (Dean and Milton, 2003; Morelli et al., 2014). Geerts and Pauw (2011) demonstrated a two-fold decline in pollination rate of an ornithophilous shrub, *Erica perspicua*, in road verge fynbos of the southwestern Cape, demonstrating the indirect effects of roads on sunbird pollination through changes in bird behaviour. However, the study was conducted along a minor road that cuts through the Kogelberg Biosphere Reserve where large areas of contiguous fynbos vegetation still occur next to the road. In transformed landscapes, where the conservation value of road verge habitats is potentially highest, road effects on bird behaviour—and thus bird pollination—might be complemented by impacts of large-scale habitat modification.

Much of the land in the CFR has been transformed by agriculture, commercial forestry, urban development, and alien plant invasions. Rouget et al. (2003) found that, collectively, these transforming agents have usurped ca. 30% of the land in the CFR, and predicted that at least 30% of the remaining natural vegetation could be transformed by the year 2023. This large-scale change in land cover has led to extensive fragmentation of landscapes in the region. Birds respond to habitat modification and fragmentation in many different ways and, as is the case with roads, the response is often guild- or species-specific (Lindenmayer et al., 2003; Luck and Daily, 2003). Nevertheless, birds in general have been shown to be more sensitive to habitat quality than other factors, including patch size, shape, and location, in fynbos fragments (Dures and Cumming, 2010). Around Cape Town, urbanization reduces the functional diversity of the ecologically important nectarivorous bird guild, with only the short-billed southern double-collared sunbird occurring more than 1 km into the city (Pauw and Louw, 2012). Armstrong et al. (1996) documented a change in avian community composition from predominantly fynbos species to one dominated by species typical of riparian woodlands after fynbos vegetation had been replaced by pine plantations. They found that specialist nectarivorous sugarbirds were present only infrequently in these artificial forests. Similarly, in Mpumalanga, species diversity of grassland birds was significantly reduced where grasslands were converted to commercial plantations, whereas woodland birds benefited from this change (Allan et al., 1997). Although these studies dealt with plantations, comparable shifts in avian assemblages may be expected where habitats are altered through alien tree invasions (Richardson and Van Wilgen, 2004). Indeed, Dures and Cumming (2010) showed that the presence of invasive *Acacia saligna* was a significant driver of reduced species richness in avian communities, while Mangachena & Geerts (2017) found species-depauperate bird assemblages in riparian habitats that were invaded by *Eucalyptus camaldunensis*. Of particular concern in the latter study was the absence of nectarivores from invaded sites.

Given that both roads and large-scale habitat modification have been demonstrated to have significant impacts on the behaviour of birds, how might these two disturbances combine to affect the pollination of ornithophilous plants in road verges? Here we investigate the pollination of a bird-pollinated shrub, *Erica glandulosa*, in road verges of a fragmented and highly transformed rural landscape. We test for road impacts on pollination by comparing number of ruptured anther rings (Geerts and Pauw, 2011) of *E. glandulosa* flowers in fynbos fragments at different distances from the road. Additionally, we test whether the type of land cover next to road verges influence the number of ruptured anther rings of *E. glandulosa* flowers.

## 2. Methodology

### 2.1 Study area

The study took place in the Eastern Cape Province, South Africa, at the southeastern end of the Cape Floristic Region (CFR). We assessed pollination rates of *Erica glandulosa* along 22 km of a four-lane divided highway, the N2 national road, between the western outskirts of Port Elizabeth (–33.948°, 25.458°) and the Van Stadens area (–33.910°, 25.229°). The speed limit along the road is 120 km h^−1^. Fynbos vegetation—a Mediterranean-type shrubland typically dominated by members of the Ericaceae, Proteaceae and Restionaceae—occurs in the road verges, including copious stands of *E. glandulosa*. Road verge width is consistent throughout at ca. 12–15 m, with the first 2 m next to the road regularly mowed. The landscape surrounding this stretch of road is rural, but largely transformed and fragmented. It comprises smaller areas of fynbos, pastures and rangelands for dairy and cattle farming, and dense stands of invasive alien trees, especially *Acacia mearnsii* and *A. saligna*, but also *Eucalyptus* and *Pinus* species. The fynbos fragments along the road range in area from ca. 25 ha to 350 ha and are all bifurcated by the road. *Erica glandulosa* populations are restricted to road verges where these abut pastures/rangelands or alien thickets; where road verges border fynbos, *E. glandulosa* stands can be found at distances more than 100 m from the road.

### 2.2 Study species

*Erica glandulosa* Thunb. (Figure 1 a) is an upright, rounded shrub growing up to 1.5 m tall (Oliver, 2012; Schumann et al., 1992). Four subspecies are recognised (Oliver and Oliver, 2005), with the typical subspecies being the subject of this study (referred to as *E. glandulosa* throughout for the sake of brevity). It occurs on well-drained, acidic sands along the coastal plains and mountains of the southern and southeastern CFR from Mossel Bay to Port Elizabeth, and is a common component of road verge fynbos between Humansdorp and Port Elizabeth (Baker and Oliver, 1967). Flowers are borne in dense, spike-like synflorescenses (Figure 1 b) at the branch tips (Oliver and Oliver, 2005; Schumann et al., 1992) and are produced year-round (Oliver, 2012). The corolla is tubular and curved, 18-26 mm long, with a pink to orange colour, or occasionally white (Baker and Oliver, 1967). Plants encountered during this study consistently featured pink flowers as is typical of populations between Humansdorp and Port Elizabeth. Plants do not resprout and rely instead on reproduction from seed to re-establish after disturbance (Hitchcock, 2013). The flowers produce copious amounts of nectar (B.A. Grobler pers. obs., 2016) that, together with their pink colour, attract sunbirds (Hitchcock, 2013). Pollen is deposited onto the beak of the bird as it disturbs the anthers during the process of nectar extraction.

**Figure 1.**
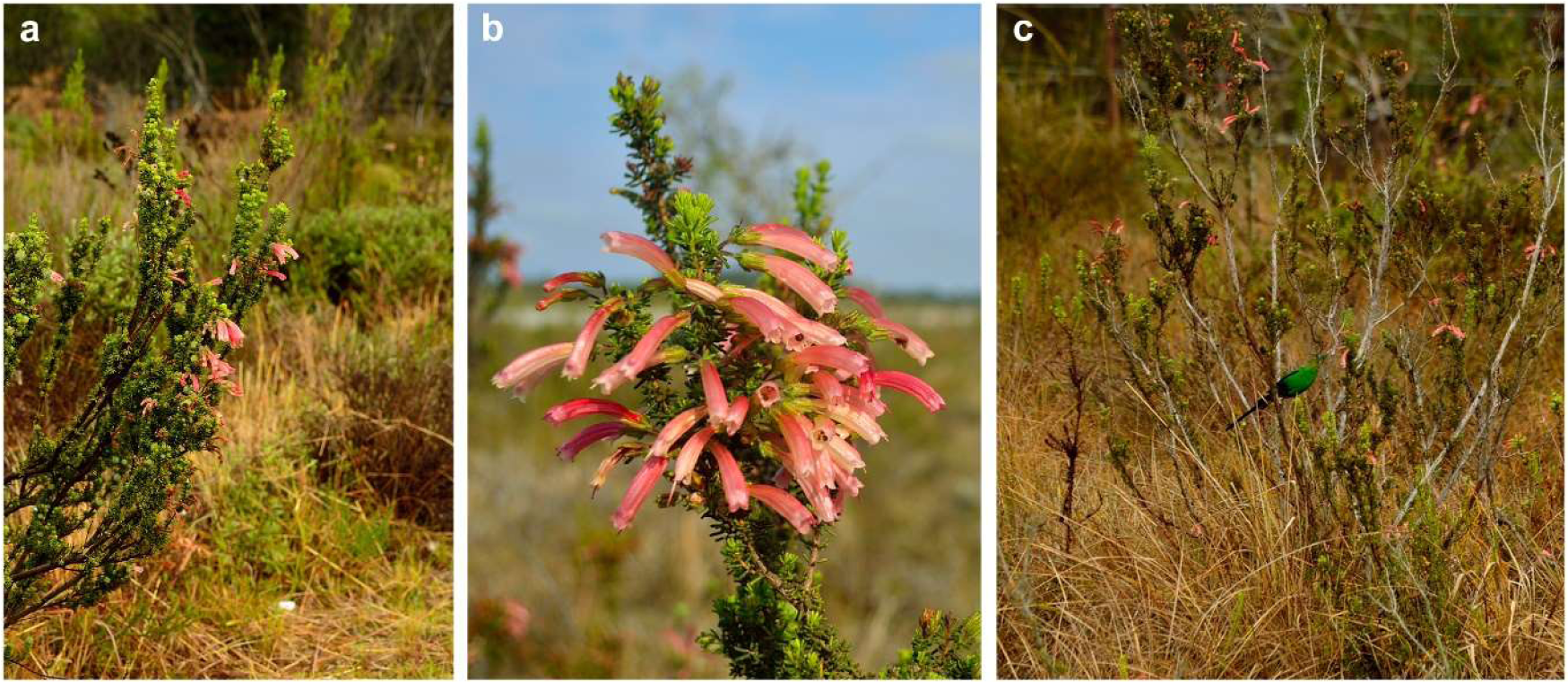
(a) *Erica glandulosa* growing in road verge fynbos along the N2 national road. (b) The pink, tubular flowers of ornithophilous *E. glandulosa* arranged in a dense, spike-like synflorescens. (c) A male malachite sunbird (*Nectarinia famosa*) visiting flowers of *E. glandulosa* in road verge fynbos.

### 2.3 Road impact on pollination

We used a technique developed by Geerts and Pauw (2011) to determine pollination rates of *E. glandulosa*. This technique employs anther ring status (intact/ruptured) as a proxy for sunbird visitation and stigma pollen receipt and is an efficient means of assessing pollination rate in ornithophilous *Erica* species (e.g., Turner et al., 2012). We established 10 m × 10 m plots within *E. glandulosa* stands at 0-10 m, 20-30 m and 40-50 m from the road edge (0 m denotes the point at which fynbos vegetation began) at nine sites along the road where fynbos occurred in the road verge and on adjacent land (Figure 2 a). Within each plot, five random flowers from each of 16 randomly selected plants were scored for ruptured anther rings and robbing. Wilted flowers were excluded as their anthers could have parted due to the withering process rather than sunbird visitation (Geerts and Pauw, 2011). Cape honeybees (*Apis mellifera* subsp. *capensis*) were primarily responsible for nectar robbing, which was indicated by the presence of a small incision towards the base of a flower (Geerts and Pauw, 2011). Plant density of *E. glandulosa* was also recorded for each plot. While sunbirds are known to preferentially visit pink flowers in colour dimorphic *Erica* species (Heystek et al., 2014), no white-flowered colour morphs of *E. glandulosa* were present in any of our plots, and flower colour was not included as a factor. In total, 720 flowers from 144 plants in nine plots were assessed for each distance interval.

**Figure 2.**
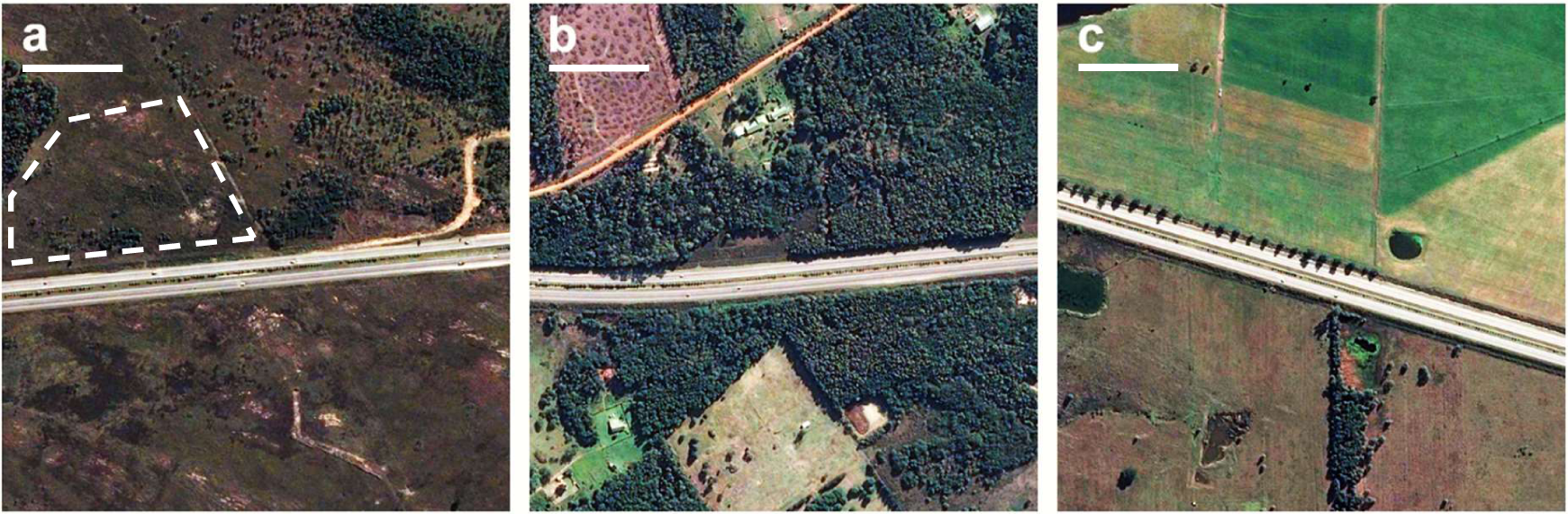
Examples of land-cover types next to road verge populations of *Erica glandulosa* along the N2 national road: (a) fynbos; (b) alien thickets; and (c) pastures (top) and rangelands (bottom). The highlighted area in (a) shows an example of sparsely invaded fynbos. Scale bars = 100 m. Images: Google, CNES/Astrium.

### 2.4 Land-cover impact on pollination

We used the same methodology described above to investigate the effect of adjacent land cover on pollination of road verge *E. glandulosa* populations. For this part of the study, 10 m × 10 m plots were located in road verge stands of *E. glandulosa*, starting at the road edge. Plots were positioned adjacent to the three major land-cover types present in the study area, namely fynbos, alien thickets, and pastures/rangelands (Figure 2). Twelve plots (960 flowers from 192 plants) were assessed adjacent to each land-cover type. Fynbos bordering road verges was not always free of alien vegetation, but plots were only located at sites where invasions were relatively sparse and localised, and where the general structure and composition of the vegetation remained intact (Figure 2 a).

### 2.5 Data analysis

All data analyses and statistical tests were implemented using R 3.6.1 (R Core Team, 2019). We used generalised linear mixed models (GLMMs), applied with the ‘glmer’ function in the ‘lme4’ R package (Bates et al., 2015), to analyse the response of pollination to road distance and adjacent land cover. These models had a binomial error distribution with a logit link function, and estimated parameters by maximum likelihood (Laplace approximation). Road distance and adjacent land cover were included as fixed effects in their respective models, while nectar robbing and plant density were included as fixed effects in both models. Data for plant density were log-scaled and mean-centred to reduce the effects of extreme values and to negate problems associated with collinearity (Zuur et al., 2009). Random effects were specified in the models to account for autocorrelation inherent in the data due to the nested sampling design. For each model, we used the maximal random effects structure corresponding to the specific nested sampling design (Barr et al., 2013). For the road impact GLMM, random effects were individual plants, plots, and sampling sites, nested in that ascending order, while individual plants nested in plots (plots and sites were synonymous in this sampling protocol) were included as random effects in the land-cover impact GLMM.

Significance tests for all fixed effects in the GLMMs were executed by means of likelihood-ratio tests (Type II sums of squares) using the ‘mixed’ function in the ‘afex’ R package (Singmann et al., 2017). This method fits the full model and various restricted models in which the parameters corresponding to each of the specified fixed effects are removed. Comparisons between the full model and the restricted models are then used to produce the test statistics. Tukey HSD *post-hoc* tests with Bonferroni correction for multiple comparisons, implemented via the ‘glht’ function in the ‘multcomp’ R package (Hothorn et al., 2008), were used to test for differences in pollination between the three distance plots and between the three land-cover types.

## 3. Results

### 3.1 Notes on floral visitors

Two bird species were seen visiting the flowers of *Erica glandulosa*, namely the malachite sunbird (*Nectarinia famosa*) and the southern double-collared sunbird (*Cinnyris chalybea*). Malachite sunbirds (Figure 1 c) were the most common visitors and were observed at all fynbos sites and most alien thicket sites (75%). Southern double-collared sunbirds were observed at only three fynbos sites (25%) and three alien thicket sites (25%). No birds were seen at pasture sites. Cape honeybees (*Apis mellifera* subsp. *capensis*) also frequented *E. glandulosa* flowers and were seen at most sites, although they were most abundant in road verges next to alien thickets.

### 3.2 Road impact on pollination

Road distance had an effect on bird pollination (Table 1; GLMM likelihood-ratio test, χ^2^ = 62.76, *P* < 0.001), with lower pollination rates next to the road than farther away from the road (Figure 3; Tukey HSD *post hoc, P* < 0.001). Pollination rates were marginally higher at 40–50 m compared to 20–30 m (Tukey HSD *post hoc, P* = 0.10). On average, pollination of *E. glandulosa* within 10 m of the road declined by 35% compared to pollination of plants at 20–30 m from the road, and by 41% compared to plants at 40–50 m from the road (Figure 3). Flower robbing rate and *E. glandulosa* plant density varied between plots at different distances from the road (Table 2). However, we found no evidence of these two factors affecting pollination (Table 1).

**Table 1:** Significance tests for the effects of road distance, adjacent land cover, nectar robbing, and plant density on percentage of disturbed anther rings (pollination rate) in *Erica glandulosa* by means of generalised linear mixed models likelihood-ratio tests.

| | *d.f.* | $\chi^2$ | *P* |
| --- | --- | --- | --- |
| *Road impact model:* |  |  |  |
| Road distance * | 2 | 62.76 | < 0.001 |
| Nectar robbing | 1 | 0.49 | 0.48 |
| Plant density † | 1 | 0.88 | 0.35 |
| *Land-cover impact model:* |  |  |  |
| Adjacent land cover * | 2 | 8.03 | 0.02 |
| Nectar robbing | 1 | 0.66 | 0.42 |
| Plant density † | 1 | 0.14 | 0.71 |

\* Significant effect at $\alpha = 0.05$
† Log-scaled and mean-centred

**Table 2:** Summary of *Erica glandulosa* flower robbing rate and plant density at different distances from the road and next to different land-cover types.

|  | Mean robbing rate (%)<br>± SE (*n* flowers) | Mean plant density (m<sup>-2</sup>) ±<br>SE (*n* plots) |
| --- | --- | --- |
| *Road distance:* |  |  |
| 0–10 m | 10.14 ± 4.42 (720) | 0.97 ± 0.14 (9) |
| 20–30 m | 12.08 ± 5.19 (720) | 0.67 ± 0.11 (9) |
| 40–50 m | 15.14 ± 5.02 (720) | 0.44 ± 0.07 (9) |
| *Adjacent land cover:* |  |  |
| Fynbos | 9.06 ± 3.32 (960) | 1.05 ± 0.16 (12) |
| Alien thicket | 21.88 ± 5.20 (960) | 0.60 ± 0.05 (12) |
| Pasture/rangeland | 12.19 ± 2.09 (960) | 0.45 ± 0.06 (12) |

**Figure 3:**
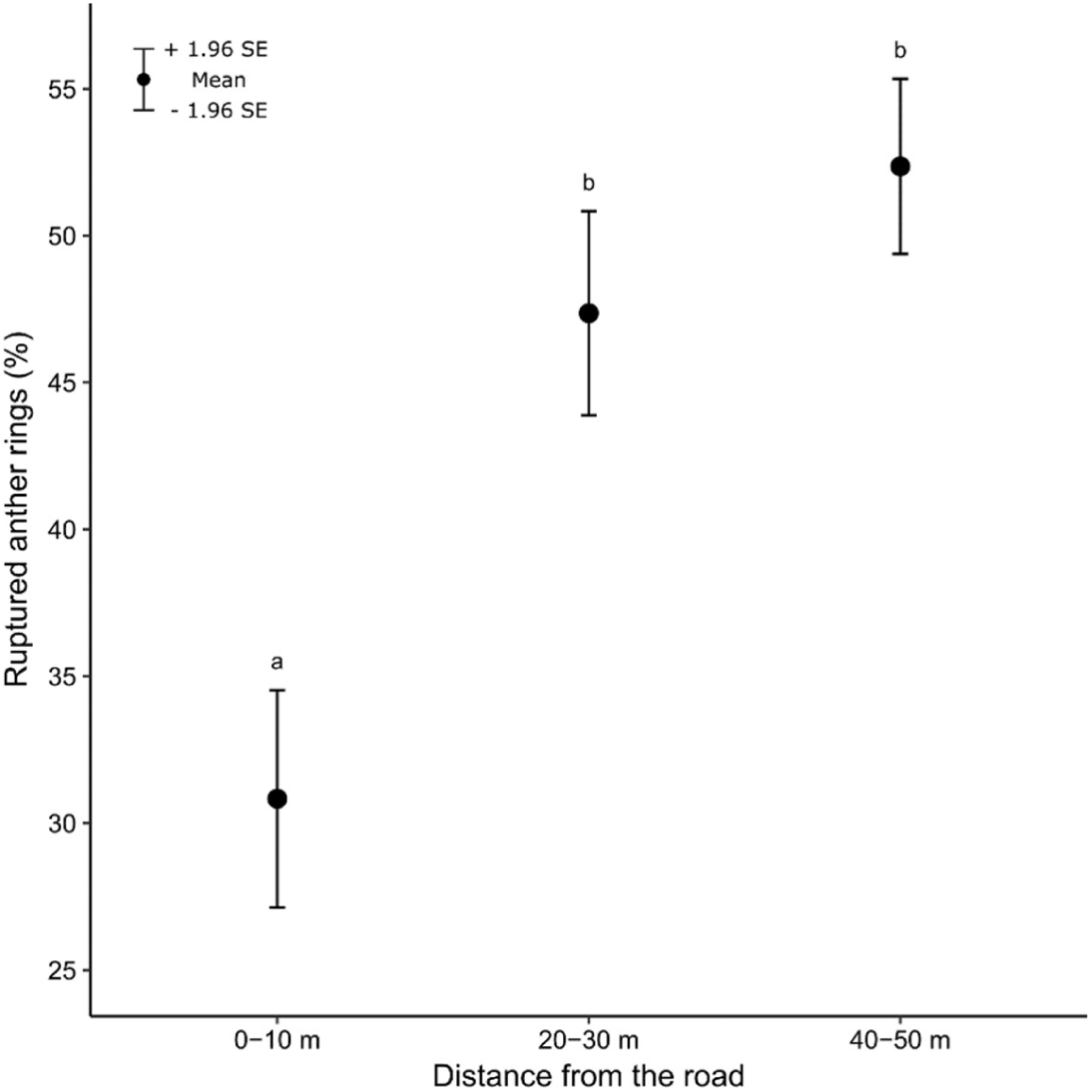
Distance from the road had a significant effect on the pollination rate of ornithophilous *Erica glandulosa* (GLMM likelihood-ratio test, χ^2^ = 62.76, *P* < 0.001). Error bars indicate ±95% confidence intervals. Different letters above bars indicate a significant difference between means (Tukey HSD *post hoc, P* < 0.001).

### 3.3 Adjacent land-cover impact on pollination

The type of land cover next to road verges impacted on bird pollination (Table 1; GLMM likelihood-ratio test, χ^2^ = 8.03, *P* = 0.02), with higher pollination rates adjacent to fynbos than next to either alien thickets or pastures/rangelands (Figure 4; Tukey HSD *post hoc, P* < 0.05). In comparison with those next to fynbos, road verge populations of *E. glandulosa* abutting alien thickets exhibited an average decline of 24% in pollination rate, while average pollination rate in populations adjacent to pastures/rangelands were 32% lower compared to those bordering fynbos (Figure 4). We found no evidence of pollination rates differing between road verge *E. glandulosa* next to alien thickets and pastures/rangelands. Flower robbing rate and *E. glandulosa* plant density varied between plots adjacent to different land-cover types (Table 2), but we could not detect an impact of these factors on pollination (Table 1).

**Figure 4:**
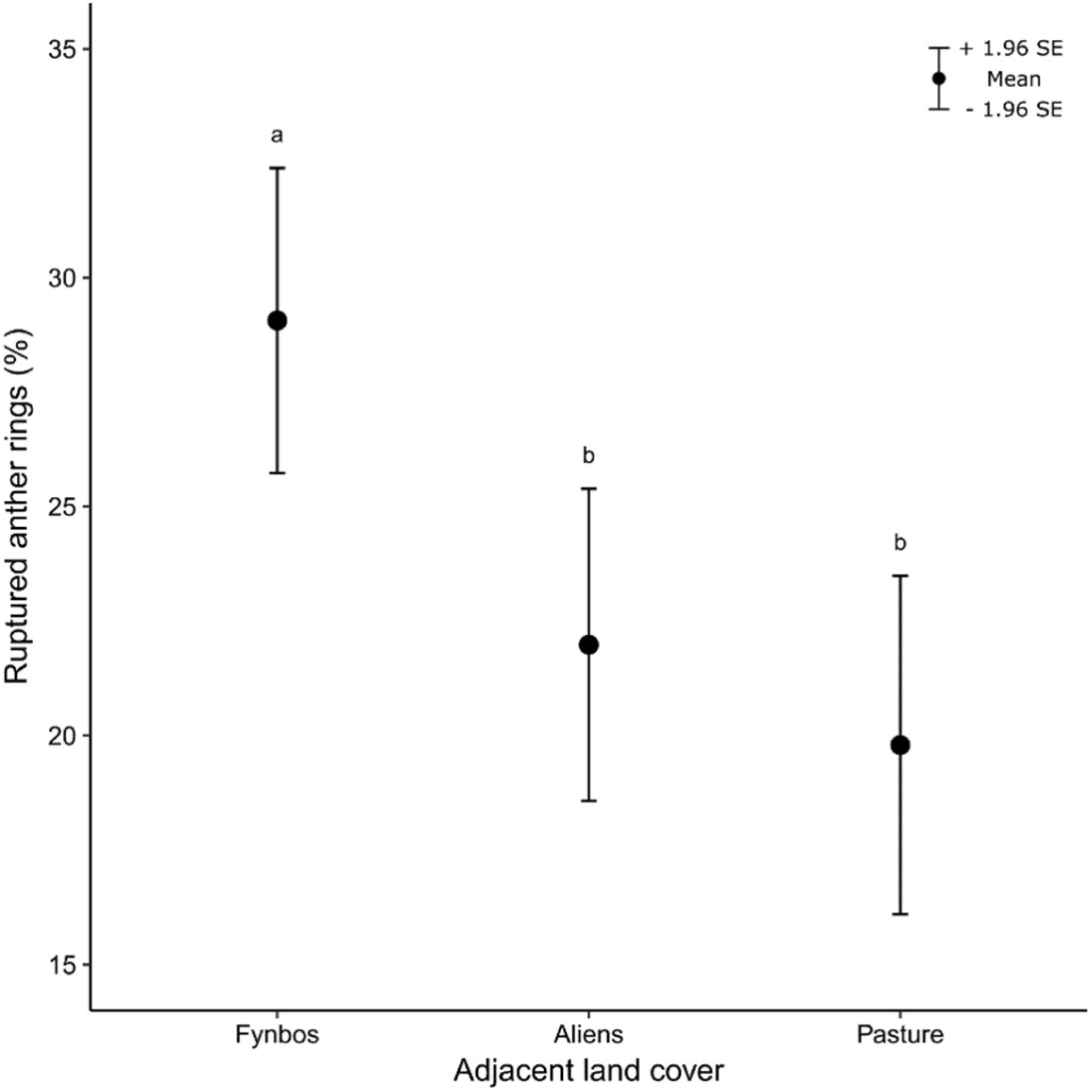
Adjacent land cover has a significant effect on the pollination of ornithophilous *Erica glandulosa* in road verge fynbos (GLMM likelihood-ratio test, χ^2^ = 8.03, *P* = 0.02). Error bars indicate ±95% confidence intervals. Different letters above bars indicate a significant difference between means (Tukey HSD *post hoc, P* < 0.05).

## 4. Discussion

### 4.1 Road impact on bird pollination

The results of this study indicate that a road impacts negatively on bird pollination in fynbos fragments of a transformed rural landscape in the southeastern CFR. On average, pollination of *E. glandulosa* within 30 m of the road declined by ca. 35–40% compared to plants farther away. These findings are consistent with those of Geerts and Pauw (2011), who showed that a two-lane tar road in the southwestern CFR caused a two-fold decline in pollination of ornithophilous *Erica perspicua* near the road. In comparison to the road investigated in the current study (a highway), the one investigated by Geerts and Pauw (2011) (a regional road) is smaller and carries less traffic at lower speeds. The impact of this regional road on pollination was evident up to 10 m away, but larger, busier roads are expected to have greater impacts on bird behaviour reaching farther from the road (Forman et al., 2002; Reijnen et al., 1996; Van der Zande et al., 1980). Although the evidence is not very strong, the results of our study support this notion as pollination rates were marginally higher at 40–50 m from the road compared to 20–30 m (Figure 3), suggesting that birds visit flowers beyond 30 m from the road more often. Nevertheless, the road impact on bird pollination remains most significant within 10 m of the road, after which it appears to diminish.

While we did not explicitly investigate the cause of reduced bird pollination nearer the road, disturbance of birds by traffic is the most likely reason. In their review of road effects on avian populations, Kociolek et al. (2011) identified traffic noise as the most significant indirect road impact that prompts road avoidance in birds. Through an experiment where synthetic traffic noise was applied to a roadless area, McClure et al. (2013) showed that, in the absence of other confounding road effects (e.g. chemical pollution, roadkill, and visual disturbance), traffic noise could reduce bird abundance by over a quarter, and elicited an almost complete avoidance in certain species. Traffic noise makes it difficult for birds to detect song by conspecifics, which in turn hinders the attraction of mates (Pohlman et al., 2009; Rheindt, 2003), the establishment and maintenance of territories (Habib et al., 2006; Mockford and Marshall, 2009), the detection of predators (Francis et al., 2009; Slabbekoorn and Ripmeester, 2008), and general communication with flock members or offspring (Leonard and Horn, 2005; Lohr et al., 2003). Higher levels of traffic have been linked to lower population densities near roads in several bird species, with average traffic densities above 30,000 vehicles per day affecting up to 55% of species in an area (Reijnen and Foppen, 2006). In the Netherlands, for a road with an average traffic density of 50,000 vehicles per day, this effect is estimated to reach up to 365 m from the road in woodlands, and up to 560 m in grasslands (Reijnen and Foppen, 2006). In the United States, this effect on grassland birds along a highway carrying more than 30,000 vehicles per day was estimated at 1,200 m (Forman et al., 2002). In comparison to these effect-distances, the 10–30 m effect distance on bird pollination revealed in our study is relatively minor.

### 4.2 Impact of landscape context on bird pollination

Bird pollination in road verges is not only affected by road disturbance; the type of land cover that occurs next to the verge also impacts on pollination (Table 1), with plants in verges neighbouring transformed land experiencing a significant reduction in pollination (Figure 4). Since McIntyre and Hobbs (1999) proposed their conceptual model of habitat fragmentation, several studies, including those investigating avian communities (Lindenmayer et al., 2003; Palmer et al., 2008; Zurita and Bellocq, 2012), have shown that a simple binary classification of habitat versus non-habitat is inappropriate for interpreting the response of organisms to habitat fragmentation and modification. Rather, viewing habitat modification as a continuum or gradient allows for a more suitable interpretation of habitat use in transformed landscapes. Based on this premise, it is improbable that modified habitats, such as the alien thickets and pastures/rangelands considered in the current study, will entirely exclude birds and so act as an impervious barrier to prevent them from accessing resources that occur in adjacent road verges. For example, a study by Forman et al. (2002) revealed no significant effect of adjacent land use (expressed as percent built area) on the presence or breeding behaviour of birds in habitat fragments along roads, although road disturbance (distance from the road) did have a significant impact. For this same reason, it is also unreasonable to expect a complete elimination of ornithophily in road verges abutting transformed land: as the results of our study show, on average, ca. 20% of *E. glandulosa* flowers in verges neighbouring transformed land are still visited by sunbirds (Figure 4).

Although habitat modification did not preclude sunbird visitation in road verges, it did reduce the number of flowers that were pollinated. Compared to those bordering fynbos, *E. glandulosa* abutting alien thickets and pastures/rangelands exhibited an average decline of 24–32% in pollination rate. In the southwestern CFR, invasive *Acacia* thickets support a large subset of the regional avifauna (Rogers and Chown, 2014). The bird communities in these thickets have densities and species richness comparable to those found in some natural sites, but, importantly, they typically lack nectarivores. Similarly, the replacement of indigenous fynbos vegetation by pines (Armstrong et al., 1996) and eucalypts (Mangachena and Geerts, 2017) also results in the displacement of nectarivores. Increased landscape heterogeneity (i.e., habitat transformation) is associated with reduced species richness and diversity of nectarivorous birds in southern Mexico (Pineda-Diez de Bonilla et al., 2012), while this same guild prefers natural forest habitats to pastures and plantations in transformed landscapes of southeastern Australia (Hsu et al., 2010). In Belize, the expansion of maintained pastures have led to a significant reduction in species richness of bird communities, and the absence of nectar-feeding birds is conspicuous in these depauperate assemblages (Saab and Petit, 1992). The development of a golf estate in the southwestern Cape, which largely replaced fynbos habitats with built-up areas and golf greens similar in composition and structure to maintained pastures, likewise resulted in reduced abundance of nectarivores (Fox and Hockey, 2007). This bird guild thus appears to be particularly sensitive to habitat modification, including the transformation of natural habitat to land-cover types so prevalent in the landscape surrounding road verges investigated in our study, as well as elsewhere in the CFR. The conversion of fynbos to alien thickets and pastures/rangelands drastically alters the composition and structure of the vegetation and reduces the diversity of native plants (Holmes and Cowling, 1997; Le Roux, 1988; Richardson et al., 1989). This causes a reduction in food resources—in terms of both abundance and diversity—which has been linked to lower abundances of specialist nectar-feeding birds (Blake and Loiselle, 2001; Chalmandrier et al., 2013; Driscoll, 1977; Geerts et al., 2011; Gray et al., 2007). It seems likely then that lower abundances of nectarivorous birds in these modified habitats is the most probable cause of reduced pollination of *E. glandulosa* in verges next to transformed land. However, this hypothesis remains to be tested.

## 5. Conclusion

The results of this study have demonstrated the impacts of road disturbance and landscape context on the pollination of ornithophilous road verge plants. To determine whether such a disrupted mutualism has important ecological implications for specific plant species, it is important to consider whether seed set is pollen limited, and whether population growth is seed limited (Anderson et al., 2011; Pauw and Bond, 2011). However, declines in pollination and subsequent pollen limitation have been shown to be the most proximate cause of reduced reproductive success—in terms of lower fruit- or seed-set—in plant populations in fragmented landscapes (Aguilar et al., 2006). Thus, even though reproductive output was not quantified in the current study, the demonstrated reduction in pollination of road verge plant populations is expected to have a negative effect on the fecundity of plants in verges. This potentially calls into question the suitability of road verges as refugia for bird-pollinated plant species in transformed landscapes. However, our results also indicate that pollination by birds is not excluded in these settings: on average, birds still visited ca. 20–30% of road verge plants. Whether this rate of pollination is sufficient to sustain populations of bird-pollinated, seed-dependent road verge plants remains to be tested.

## Competing Interests

The authors declare that they have no competing interests.

## Author Contributions

BAG designed the study, collected the data, analysed the data, prepared the figures and tables, authored drafts of the manuscript and approved the final draft. EEC reviewed drafts of the manuscript and approved the final draft.

## Acknowledgments

The South African National Research Foundation (NRF) and the South African National Roads Authority (SANRAL) is thanked for financial support. Roland Thompson from SANRAL is thanked for permission to access land under his management. We are grateful to Shandon Carvalho for his assistance in the field.

